# Base editing generates substantial off-target single nucleotide variants

**DOI:** 10.1101/480145

**Authors:** Erwei Zuo, Yidi Sun, Wu Wei, Tanglong Yuan, Wenqin Ying, Lars M. Steinmetz, Yixue Li, Hui Yang

**Author notes:** These authors contributed equally to this work (Erwei Zuo, Yidi Sun, Wu Wei, Tanglong Yuan).

## Abstract

Genome editing tools including CRISPR/Cas9 and base editors hold great promise for correcting pathogenic mutations. Unbiased genome-wide off-target effects of the editing in mammalian cells is required before clinical applications, but determination of the extent of off-target effects has been difficult due to the existence of single nucleotide polymorphisms (SNPs) in individuals. Here, we developed a method named GOTI (Genome-wide Off-target analysis by Two-cell embryo Injection) to detect off-target mutations without interference of SNPs. We applied GOTI to both the CRISPR-Cas9 and base editing (BE3) systems by editing one blastomere of the two-cell mouse embryo and then compared whole genome sequences of progeny-cell populations at E14.5 stage. Sequence analysis of edited and non-edited cell progenies showed that undesired off-target single nucleotide variants (SNVs) are rare (average 10.5) in CRISPR-edited mouse embryos, with a frequency close to the spontaneous mutation rate. By contrast, BE3 editing induced over 20-fold higher SNVs (average 283), raising the concern of using base-editing approaches for biomedical application.

---

CRISPR/Cas9 and base editors mediated genome editing methods have been developed and carried great hope for the treatment of genetic diseases induced by pathogenic mutations (1) (2) (3) (4) (5) (6). Clinical applications of CRISPR/Cas9-based genome editing or base editors require a comprehensive analysis of off-target effects to minimize risk of deleterious mutations, but a well validated method robust to genetic variants is undescribed yet to detect SNVs (7). Multiple methods have been developed to detect genome-wide gene-editing off-target activity in cells, including high-throughput genome-wide translocation sequencing (HTGTS) (8), genome-wide, unbiased identification of double-strand breaks (GUIDE-seq) (9) and circularization for in vitro reporting of cleavage effects by sequencing (CIRCLE-seq) (10). However, these approaches are not applicable to detect SNVs. Thereafter, GOTI was developed and applied to evaluate the off-target effects induced by CRISPR/Cas9 or BE3 (rAPOBEC1-nCas9-UGI), the commonly used gene editing tools (4, 11). Briefly, we performed either CRISPR/Cas9 or BE3 gene editing, together with Cre mRNA, in one blastomere of the two-cell embryos derived from Ai9 (CAG-LoxP-Stop-LoxP-tdTomato) mice (12, 13) (Fig. 1a). The progeny cells of edited and non-edited blastomeres were then sorted at E14.5 by fluorescence-activated cell sorting (FACS), based on tdTomato expression in the gene-edited cells. Whole genome sequencing (WGS) was then performed on the tdTomato^+^and tdTomato^-^ cells, separately (Fig. 1a).

**Figure 1.**
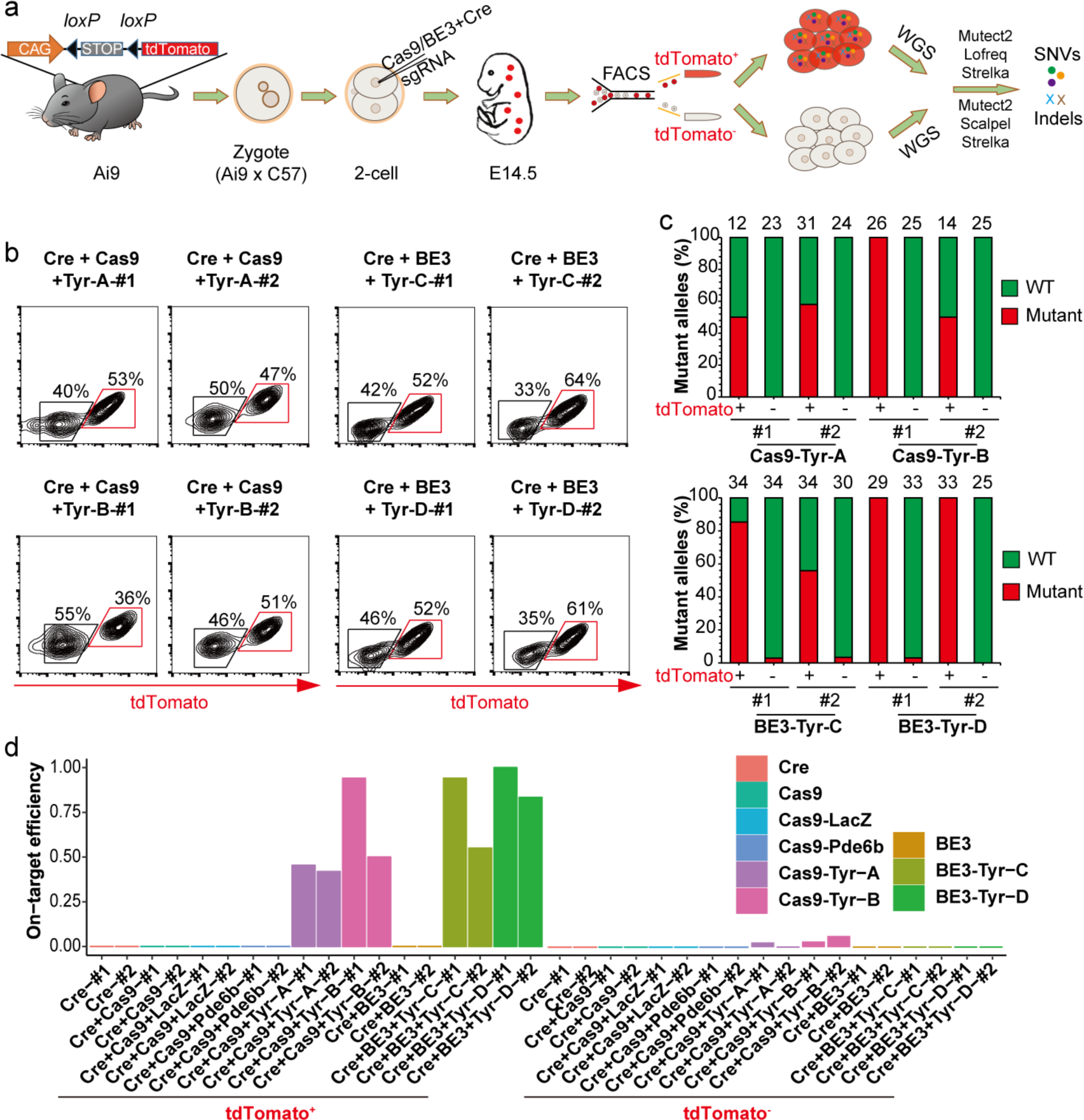
CRISPR/Cas9-or BE3-mediated gene editing in one blastomere of 2-cell embryos. **a**, Experimental design. The mixture of Cre, Cas9/BE3 and sgRNA was injected into one blastomere of a 2-cell embryo, derived from Ai9 male mice mating with wild-type female mice. The action of Cre is expected to generate a chimeric embryo with half of cells labeled with tdTomato (colored red). The tdTomato^+^ cells and tdTomato^-^ cells were isolated from the chimeric embryo at E14.5 by FACS and used for WGS respectively. The off-target SNVs and indels were identified by comparing the tdTomato^+^ cells with tdTomato^-^ cells using three variant calling algorithms as indicated (Mutect2, Lofreq and Strelka for SNVs, and Mutect2, Scalpel and Strelka for indels). SNVs and indels were indicated as colored dots and crosses. **b**, FACS analysis on E14.5 embryos treated with Cas9-Tyr-A, Cas9-Tyr-B, BE3-Tyr-C or BE3-Tyr-D. FACS analysis of uninjected embryo was shown in fig. S1b. **c**, On-target analysis of TA clone sequencing on E14.5 embryos treated with Cas9-Tyr-A, Cas9-Tyr-B, BE3-Tyr-C or BE3-Tyr-D. Number above each column, total clones analyzed. **d**, On-target efficiency for the tdTomato^+^ cells (left) and tdTomato^-^ cells (right) included in the study based on WGS. Embryos with the same treatment were represented in the same color.

To validate our approach, we demonstrated that edited cells treated with Cre and Cas9/BE3 system was efficiently separated with non-edited cells. Upon Cre-mediated recombination, about 50% of the embryo cells are expected to express tdTomato. This was verified at the 4- and 8-cell stages by fluorescence microscopic observation (fig. S1a, b) or at E14.5 by FACS analysis (Fig. 1b). Furthermore, we found that editing the coat-color gene tyrosinase *(Tyr)* with CRISPR/Cas9 by injection of one blastomere of 2-cell embryos with either one of the two sgRNAs (Cas9-Tyr-A, Cas9-Tyr-B) resulted in highly efficient on-target editing: Sanger sequencing of *Tyr* gene showed that 13 (for Cas9-Tyr-A) and 15 (for Cas9-Tyr-B) tdTomato^+^ cells individually collected from four dispersed 8-cell embryos, 85% and 80% of cells carrying *Tyr* mutated alleles, respectively. By contrast, none of tdTomato^-^ cells collected (16 for Cas9-Tyr-A and 15 for Cas9-Tyr-B) showed *Tyr* mutated alleles (fig. S1c).

In addition, we verified the editing efficiency of our approach on the targeted *Tyr* gene. For genome-wide sequencing studies using the embryo injection protocol, we included four sgRNAs for CRISPR/Cas9 editing: Cas9-Tyr-A and Cas9-Tyr-B targeting at *Tyr*, one control sgRNA targeting at *LacZ* without cleavage sites in the C57 mouse genome, and a sgRNA targeting at *Pde6b*, which has one mismatch with the C57 mouse genome and was previously reported to generate thousands of SNVs (14). We validated the cleavage efficiency of these sgRNAs (~100%) *in vitro* using DNA cleavage assays (fig. S2). In addition, we examined two sgRNAs (BE3-Tyr-C and BE3-Tyr-D) that targeted *Tyr* gene via BE3-mediated editing. As control groups for gene editing, we also included three groups of embryos injected with “Cre only”, “Cre and Cas9” and “Cre and BE3”. After injection of different mixtures of CRISPR/Cas9 or BE3, and Cre mRNAs as well as sgRNAs into one blastomere of two-cell embryos, we found no impairment of embryonic development, as indicated by the normal blastocyst rate and survival rate (fig. S3a-b). By Sanger sequencing, all the examined blastocysts and E14.5 fetuses derived from Cas9-Tyr-A, Cas9-Tyr-B, BE3-Tyr-C and BE3-Tyr-D carried *Tyr* mutations (fig. S3c). To evaluate target-editing efficiencies for the *Tyr* gene, we FACS-sorted about four million tdTomato^+^ and four million tdTomato^-^ cells from one E14.5 embryo treated with either Cas9-Tyr-A or Cas9-Tyr-B, and performed TA clone sequencing. The results from one experiment showed that tdTomato^+^ cells carried 50% and 100% mutated alleles (with 12 and 26 clones were examined) for Cas9-Tyr-A and Cas9-Tyr-B targeting, respectively (Fig. 1c). In the second repeated experiments, tdTomato^+^ cells carried 58% and 50% mutated alleles (with 31 and 14 clones examined) respectively for Cas9-Tyr-A and Cas9-Tyr-B targeting (Fig. 1c). By contrast, tdTomato^-^ cells collected from 14.5 embryos carried no *Tyr* mutated alleles (Fig. 1c). Similarly, BE3 editing demonstrated high targeting efficiencies, with 71% mutant alleles on BE3-Tyr-C and 100% on BE3-Tyr-D, and the corresponding tdTomato^-^ cells had only ~3% *Tyr* mutations (Fig. 1c). These results suggested that both CRISPR/Cas9 and BE3 editing yield high on-target editing efficiency in tdTomato^+^ cells, but essentially no on-target editing in tdTomato^-^ cells.

To further explore the on-target efficiency and potential genome-wide off-target effects, we performed WGS at an average depth of 47x on 36 samples from 18 E14.5 embryos in nine groups (with 2 embryos each): Cre only, Cre and Cas9, Cre and Cas9-LacZ, Cre and Cas9-Pde6b, Cre and Cas9-Tyr-A, Cre and Cas9-Tyr-B, Cre and BE3, Cre and BE3-Tyr-C, and Cre and BE3-Tyr-D (Omit “Cre” in the latter eight groups for short), among which only Cas9-Tyr-A, Cas9-Tyr-B, BE3-Tyr-C and BE3-Tyr-D groups have target sites on the C57 genome (table S1). On-target analysis for Cas9-Tyr-A and Cas9-Tyr-B groups showed an average of 56% and 72% *Tyr* mutated alleles in tdTomato^+^ cells, respectively (Fig. 1d), indicating efficient on-target DNA editing at *Tyr* gene locus. Similarly, both BE3-Tyr-C and BE3-Tyr-D demonstrated high editing efficiency (averaged 75% and 92% respectively *Tyr* mutated alleles) in tdTomato^+^ cells (Fig. 1d). We also analyzed the on-target efficiency for all the other tdTomato^+^ embryos, and found zero on-target editing for all the control embryos (Fig. 1d). However, the WGS analysis also revealed a low-level of targeted editing in tdTomato^-^ cells of Cas9-Tyr-A and Cas9-Tyr-B groups in the range of 0 - 6.3% (Fig. 1d, fig. S4 and table S2), which was probably caused by false-negative FACS sorting (known to occur at a low level). Thus we only considered variants with allele frequencies more than 10% to be reliable in our following analysis.

To assess the off-target editing effects, we analyzed the genome-wide *de novo* variants by comparing the tdTomato^+^ cells with the tdTomato^-^ cells in each embryo with three different variant calling algorithms simultaneously (14, 15), with variants defined by all three algorithms to be the true variants (Fig. 2a, fig. S5, table S3 and Methods). We found only 0 - 4 indels in embryos from all nine groups, a result further validated by Sanger sequencing (Table 1, fig. S6a and table S3). Meanwhile, in the “Cre only” embryos, we observed an average of 14 SNVs (Table 1 and table S4). For the CRISPR/Cas9-treated embryos (Cas9, Cas9-LacZ, Cas9-Pde6b, Cas9-Tyr-A, and Cas9-Tyr-B), there were an average of 12.5, 5, 0, 16 and 19 SNVs, respectively in each group (Table 1 and table S4), and no significant difference among them (Fig. 2a) or in comparison with the “Cre-only” group (Fig. 2a-b). In addition, we observed no increased SNVs in the Cas9-Pde6b edited embryo, consistent with many previous studies (16-19). All off-target SNVs detected in the CRISPR/Cas9 edited embryos were further confirmed by Sanger sequencing (fig. S6b). The SNV variants detected in the Cre- or CRISPR/Cas9-treated samples were likely caused by spontaneous mutations during genome replication in the development, since the number of variants was within the range of the simulated spontaneous mutations (fig. S7, Methods) (20). Besides, none of the samples in our study shared identical variants (fig. S8a), and no overlap was found for SNV sites predicted by Cas-OFFinder, CRISPOR and Digenome-seq putative off-target sites (fig. S8b and tables S5-7, Methods) (21, 22). We also observed no sequence similarity between the adjacent sequences of the identified SNVs with the target sites (fig. S8c). In addition, by conversely calling variants through comparing the tdTomato^-^ sample with tdTomato^+^ sample in each embryo, we found similar numbers of SNVs (fig. S9a and table S8), suggesting that CRISPR/Cas9 editing induces no off-target edits and the SNVs we observed are from spontaneous mutations.

**Figure 2.**
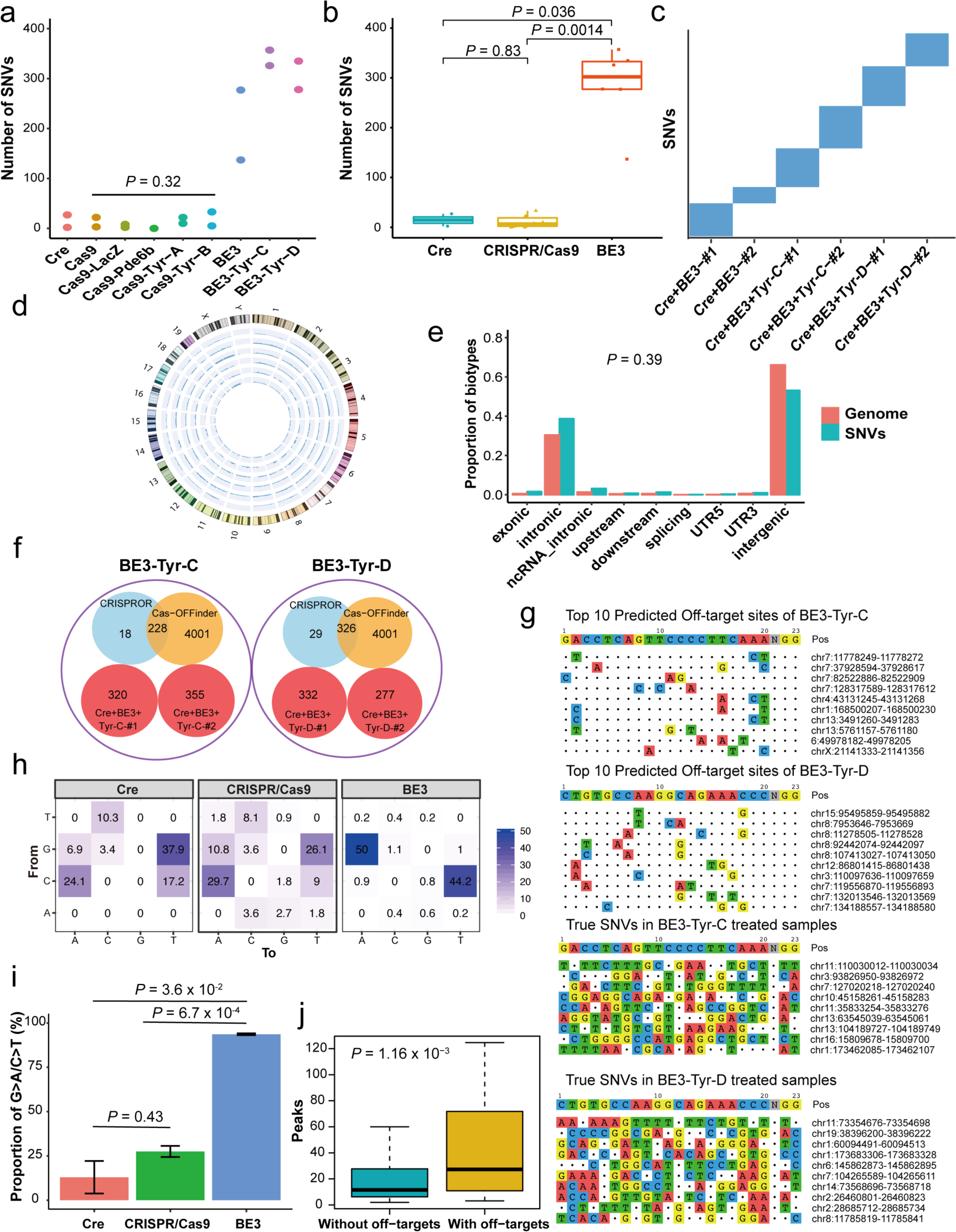
Genome-wide SNVs identified in Cre-, CRISPR/Cas9- and BE3-treated mouse embryos. **a**, The number of SNVs identified in Cre-treated, CRISPR/Cas9-treated (Cas9, Cas9-LacZ, Cas9-Pde6b, Cas9-Tyr-A, and Cas9-Tyr-B) and BE3-treated (BE3, BE3-Tyr-C, BE3-Tyr-D) groups. Each point indicated one embryo. Colors indicated different treatment conditions. *P*-value was calculated by Kruskal-Wallis test among CRISPR/Cas9-treated embryos. **b**, The comparison of the total number of detected SNVs in Cre-treated only, CRISPR/Cas9-treated and BE3-treated samples. **c**, The SNVs identified from each embryo with BE3 injected were mutually exclusive from each other. **d**, The detected SNVs were evenly distributed in the mouse genome in all the six BE3-treated samples. Embryos from inner circle to outer circle were Cre+BE3-#1, Cre+BE3-#2, Cre+BE3-Tyr-C-#1, Cre+BE3+Tyr-C-#2, Cre+BE3+Tyr-D-#1 and Cre+BE3+Tyr-D-#2, respectively. **e**, The biotypes of our detected SNVs were similar to the structure of mm10 genomes (*P* = 0.39 by Wilcoxon rank sum test). **f**, The overlap among SNVs detected from our results with predicted off-targets sites by Cas-OFFinder and CRISPOR. **g**, The sequence similarities between the targeted sequence with predicted off-target sites or our identified mutation sequences. **h**, The distribution of mutation types in Cre-, CRISPR/Cas9- and BE3-treated groups. The number in each cell indicated the proportion of a certain type of mutation among all the mutations. Deeper colors represented higher proportion of mutation types. **i**, The proportion of C>T and G>A mutations in the Cre only, CRISPR/Cas9- and BE3-treated embryos. *P* values were calculated by two-sided Wilcoxon rank sum test and *P* < 0.05 was considered as significant. **j**, The comparison of peak regions with or without the identified off-targets from a ATAC-seq dataset with GEO accession GSM2551664. *P*-values were calculated with Wilcoxon rank sum test.

**Table 1.**
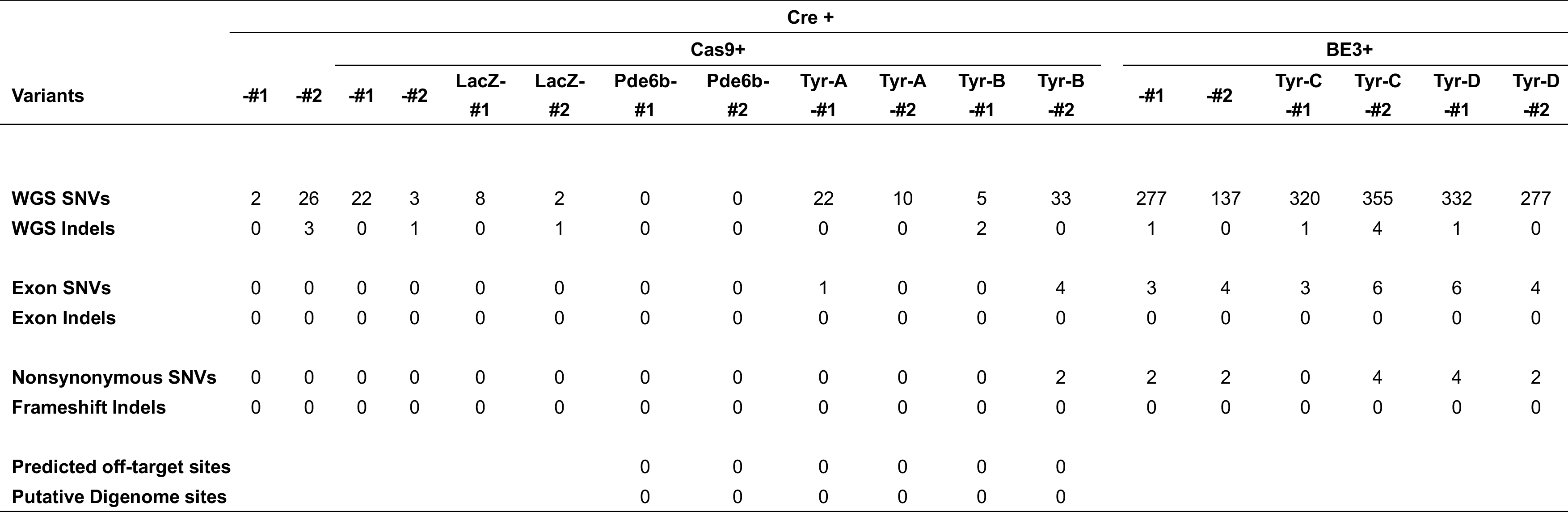
Summary of SNVs and indels identified from WGS in each embryo.

Surprisingly, in contrast with the CRISPR/Cas9 editing, we found an average of 283 SNVs in all BE3-treated embryos with or without sgRNA (BE3, BE3-Tyr-C, BE3-Tyr-D), a level of SNVs at least 20 times more than that observed in Cre-injected or CRISPR/Cas9-treated embryos (Figure 2a, b, Table 1 and fig. S5b). By conversely calling variants in each embryo (tdTomato^-^ vs tdTomato^+^), only 20 SNVs on average were detected (fig. S9b and table S9), which was presumed to be the spontaneous mutations considering the analysis on the CRISPR/Cas9-treated samples. These results indicated that the increased SNVs identified were caused by the injection of BE3.

Notably, for the off-targets detected in BE3-treated samples, none of them were shared by any of the embryos in each group (Fig. 2c), and they were randomly distributed across the genome (Fig. 2d, e, table S10). We then compared these off-target mutations with all the potential off-target sites predicted by both Cas-OFFinder and CRISPROR softwares (tables S11 and S12) (21, 22). Not surprisingly, the two prediction tools shared a large amount of off-target sites, but none of them were found in our detected SNV sites (Fig. 2f). Besides, no sequence similarity was observed between adjacent sequences of the identified SNVs and the BE3 sgRNA targeting sites, whereas the top predicted off-target sites showed sequence similarity with BE3 on-target loci (Fig. 2g). Remarkably, in spite of the uniqueness of the SNVs generated by BE3-editing, the mutation types were the same as those preferred by APOBEC1 (23, 24) (Fig. 2h and fig. S10). In fact, more than 90% of the SNVs identified in the BE3-edited cells were mutated from G to A or C to T, which was not observed in Cre- or CRISPR/Cas9-treated cells (Fig. 2h and fig. S10), indicating that mutations were not spontaneous but induced by BE3-editing. Statistical significance was also observed in the proportion of G>A and C>T among all mutations between BE3-treated cells and Cas9-(P = 6.7 x 10^-4^) or Cre-(P = 3.6 x 10^-2^) treated cells (Fig. 2i). It has been reported that DNA accessibility is related to gene editing efficiency (25). Thus we evaluated whether the identified off-target sites enriched in open chromatin regions by checking the ATAC-seq datasets derived from embryonic mouse cells in the Cistrome database (26-28). In fact, off-target sites were significantly enriched in the regions with higher accessibilities in a dataset from E8.5 embryos with a mixed C57BL6/DBA2 background (Fig. 2j) and other four high quality datasets in Cistrome database. (fig. S11).

Among 1698 SNVs induced by BE3 editing in six BE3-treated embryos, 26 were located on exons (tables S13). We successfully amplified 20 of them by PCR and confirmed their presence by Sanger sequencing (Fig. 3a, fig. S12 and tables S13). Among the 26 SNVs, 14 of them lead to nonsynonymous changes in the coded proteins, and two SNVs result in early stop-codons in *Trim23* and *Aim2* genes. The *Trim23* encodes an E3 ubiquitin-protein ligase, and its dysfunction could cause muscular dystrophy (29). Previous studies reported an essential role of *Aim2* gene in innate immunity underlying the defense against viral infection (30, 31) (Fig. 3b). We also found 1 SNV in a proto-oncogene and 13 SNVs in tumor suppressors (Fig. 3c), raising serious concern about the oncogenic risk of BE3-editing.

**Figure 3.**
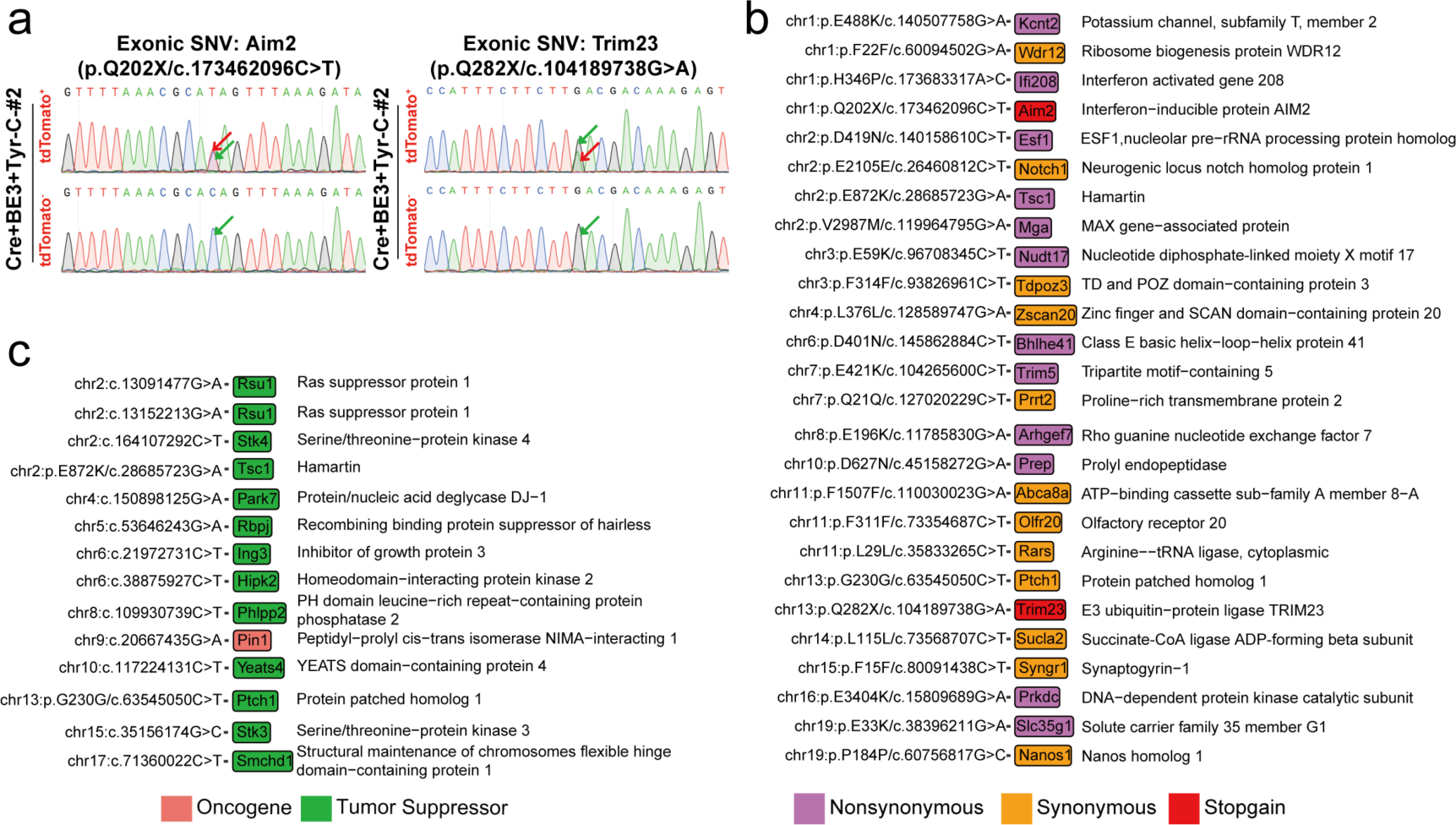
The BE3 induced off-targets could be detrimental to gene functions. **a**, Sanger sequencing validation on exonic SNVs located on *Aim2 and Trim23* genes. The mutated and WT base sequence were pointed out by red and green arrows, respectively. **b**, SNVs located on the protein coding exons. Nonsynonymous, synonymous and stop-gain SNVs were indicated by different colors. **c**, The SNVs located within the cancer associated genes. Red and green colors indicated oncogene and tumor suppressor, respectively. Primers were listed in table S16.

One of the big advantages of our approach is that rules out genetic background differences by comparing edited to non-edited cells in a single mouse. Previous studies were confounded by genetic background variations when comparing edited and non-edited mice. In fact, we also applied our analysis method to a published dataset *(15)* and found an average of ~1000 SNVs and ~100 indels between CRISPR/Cas9-edited mice and their non-edited siblings (tables S14). Based on our finding, we suggest that these differences among siblings are due to genetic variations rather than the results of CRISPR/Cas9-editing. Furthermore, when the sequences between any two different embryos in our study were compared, we observed more SNVs (3706 ± 5232) and indels (583 ± 762) (n = 18 pairs), as the embryos used in our study were not all derived from the same parents (table S15). These results illustrate that it is difficult to find a robust control in off-target analysis by comparing an edited to non-edited mouse as there is a significant amount of genetic variations between mice even when they have the same parents.

In summary, our studies demonstrate the advantages of GOTI approach using whole-genome sequencing analysis of progeny cells of sister blastomeres in studying off-target effects induced by gene-editing. We also showed that undesired off-target mutations induced by CRISPR/Cas9-mediated gene editing are rare in mouse embryos, supporting previous findings that CRISPR/Cas9-based editing *in vivo* does not cause significant SNVs and indels (15, 32–34). However, we could not exclude the big deletions or large chromosomal translocations reported by other studies (35, 36). By contrast, we found numerous *de novo* SNVs induced by BE3-editing, raising the safety concern of base editing approaches in therapeutic applications.

## AUTHOR CONTRIBUTIONS

EZ designed and performed experiments. YS and WW performed WGS analysis. TY performed PCR analysis. WY performed mouse embryo transfer. HY, YL and LS supervised the project and designed experiments. All the authors wrote the paper.

## ACKNOWLEDGMENTS

We thank Dr. Mu-ming Poo, Dang-sheng Li, Guo-liang Xu and Kevin Roy for helpful discussions and insightful comments on this manuscript. This work was supported by National Science and Technology Major Project (2017YFC1001302), CAS Strategic Priority Research Program (XDB02050007, XDA01010409), the National High-tech R&D Program (863 Program; 2015AA020307), the National Natural Science Foundation of China grants (31522037, 31500825), China Youth Thousand Talents Program (to HY), Breakthrough project of Chinese Academy of Sciences, Shanghai City Committee of science and technology project (16JC1420202, HY) and National Key Research and Development Program of China (2017YFC0908405) to WW.

## COMPETING FINANCIAL INTERESTS

The authors declare no competing financial interests.

---

**Figure S1.**
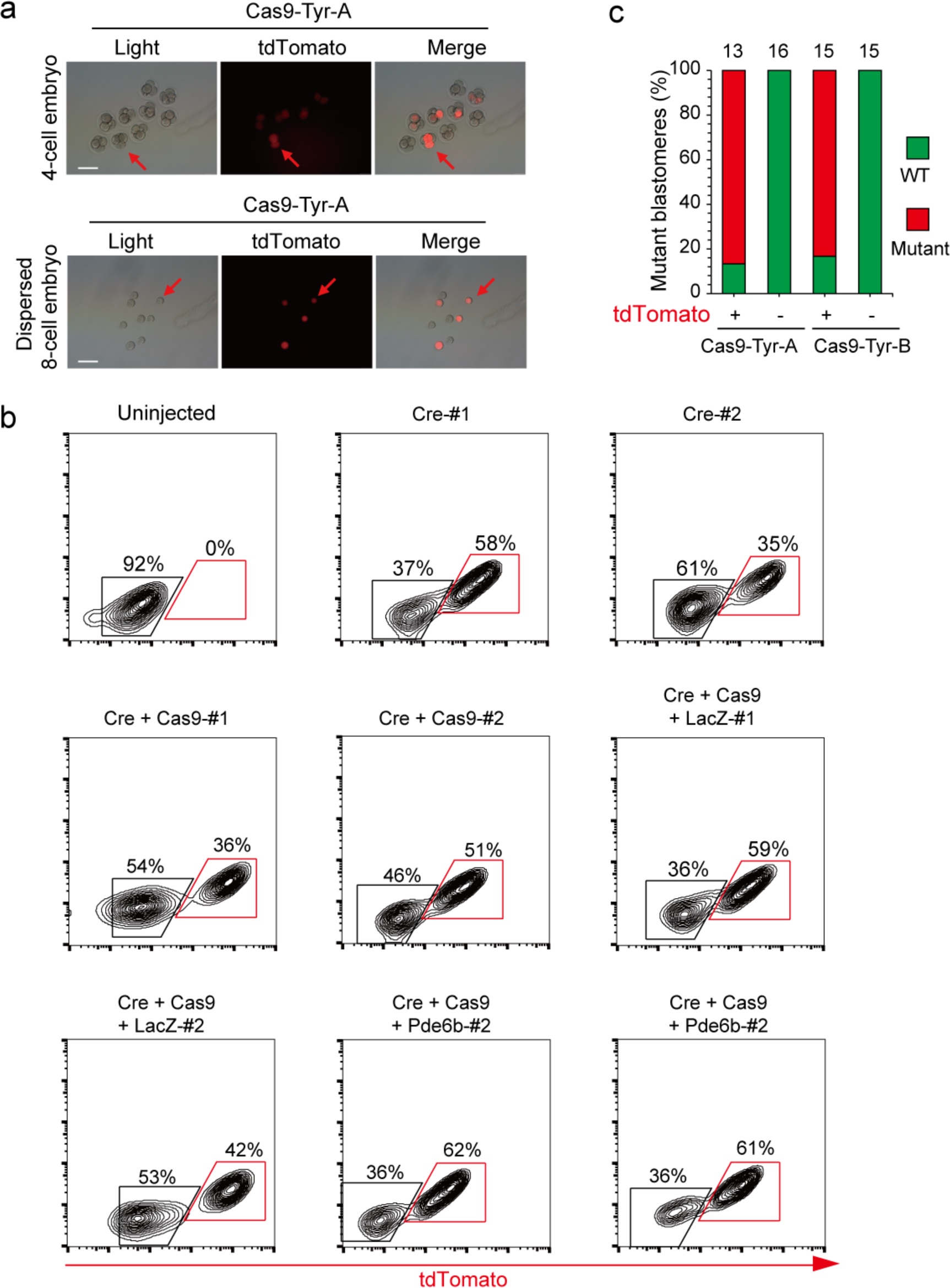
FACS analysis on E14.5 embryos treated with different mixtures. **a**, Representative images of Cas9-Tyr-A gene-targeted embryos. Top, 4-cell embryos; bottom, a dispersed 8-cell embryo. Red arrows indicate tdTomato^+^ blastomeres. Scale bar, 100μm. **b**, From left to right and top to the bottom successively: uninjected, Cre-#1, Cre-#2, Cre+Cas9-#1, Cre+Cas9-#2, Cre+Cas9+LacZ-#1, Cre+Cas9+LacZ-#2, Cre+Cas9+Pde6b-#1 and Cre+Cas9+Pde6b-#2. **c**, Genotypes of *Tyr* gene-targeted 8 cell embryos. Individual tdTomato^+^ and tdTomato^-^ blastomeres were isolated from four Cas9-Tyr-A and four Cas9-Tyr-B gene-targeted 8-cell embryos. Number, total blastomeres analyzed. WT, wild-type allele. Mutant, *Tyr* mutated allele.

**Figure S2.**
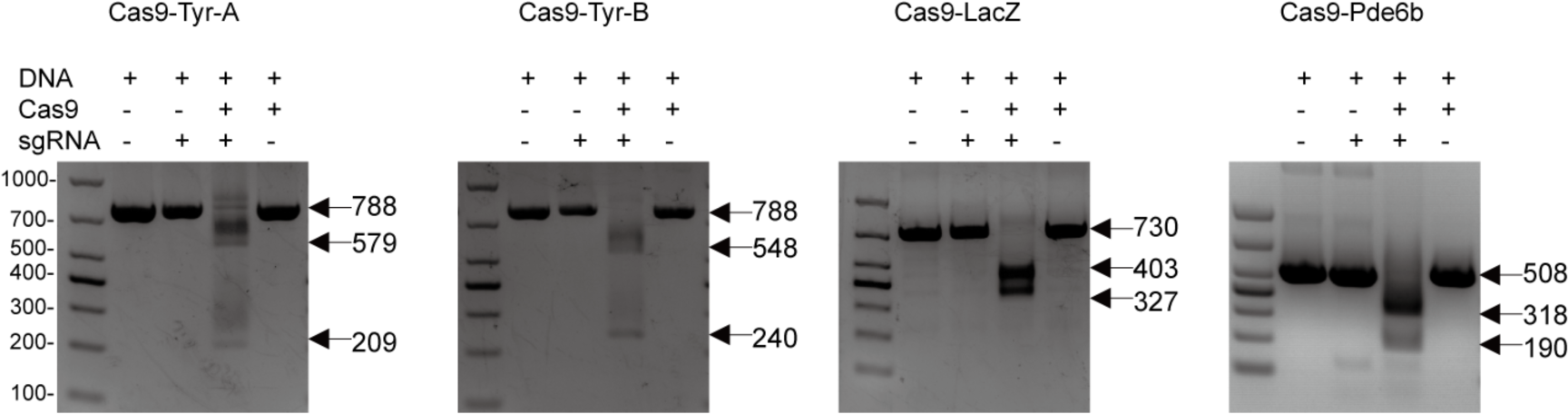
The cleavage efficiencies of sgRNAs using in vitro cleavage of DNA assay. Agarose gel electrophoresis from left to right showed the result of Cas9-Tyr-A, Cas9-Tyr-B, Cas9-LacZ and Cas9-Pde6b, respectively. The genomic region or construct flanking the sgRNA target site for each gene was PCR amplified and PCR products were incubated with Cas9 ribonucleoproteins and sgRNA for 3 h.

**Figure S3.**
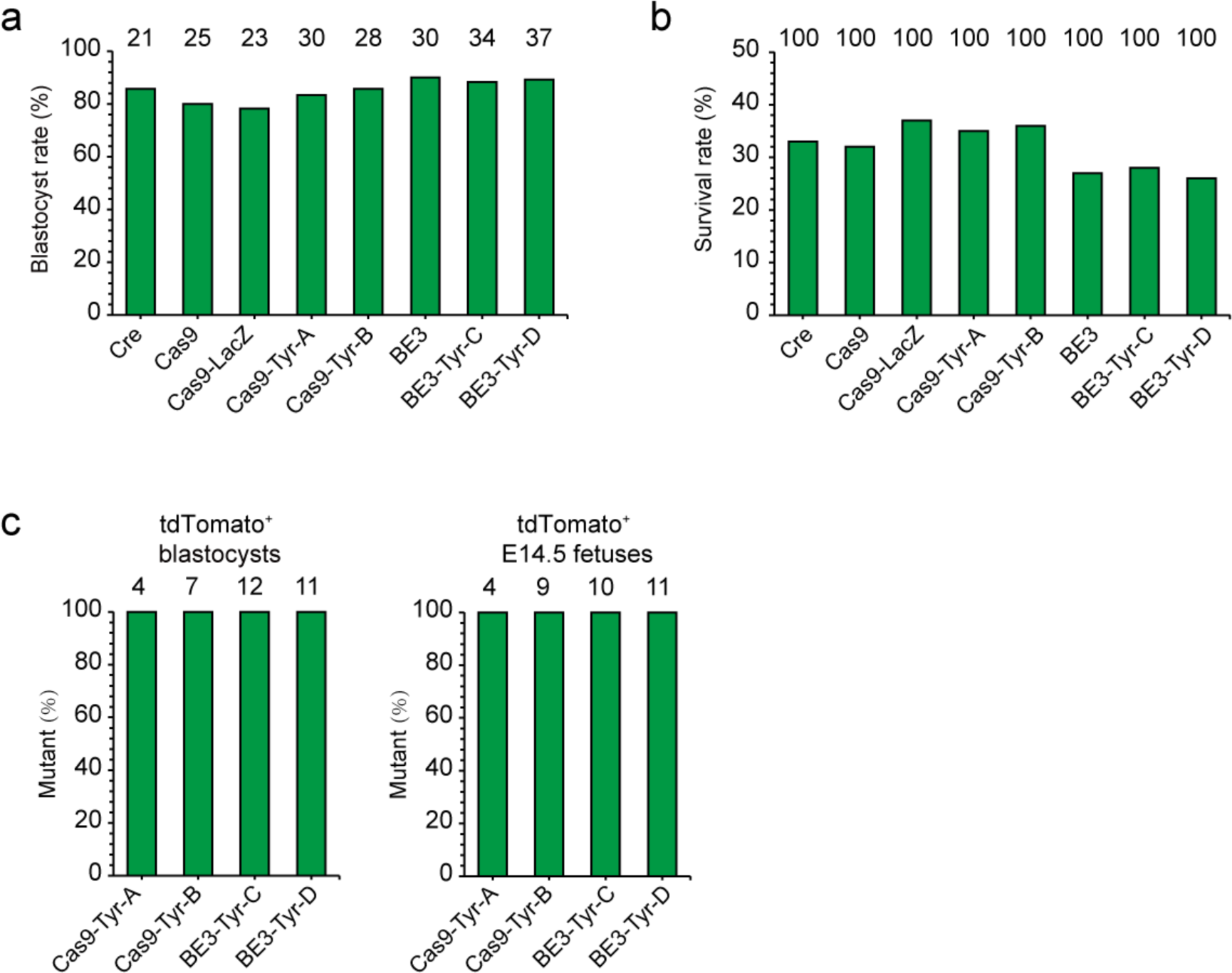
Development and genotyping of CRISPR/Cas9 and BE3-treated chimeric embryos. **a**, Percentages of tdTomato^+^ blastocysts from embryos injected with different mixtures. Number, total blastocysts or embryos counted. **b**, Survival rate at E14.5 of embryos injected with different mixtures. Number, total embryos counted or transferred. **c**, The percentage of tdTomato^+^ blastocysts and E14.5 embryos with *Tyr* mutations. Number, total blastocysts or embryos analyzed.

**Figure S4.**
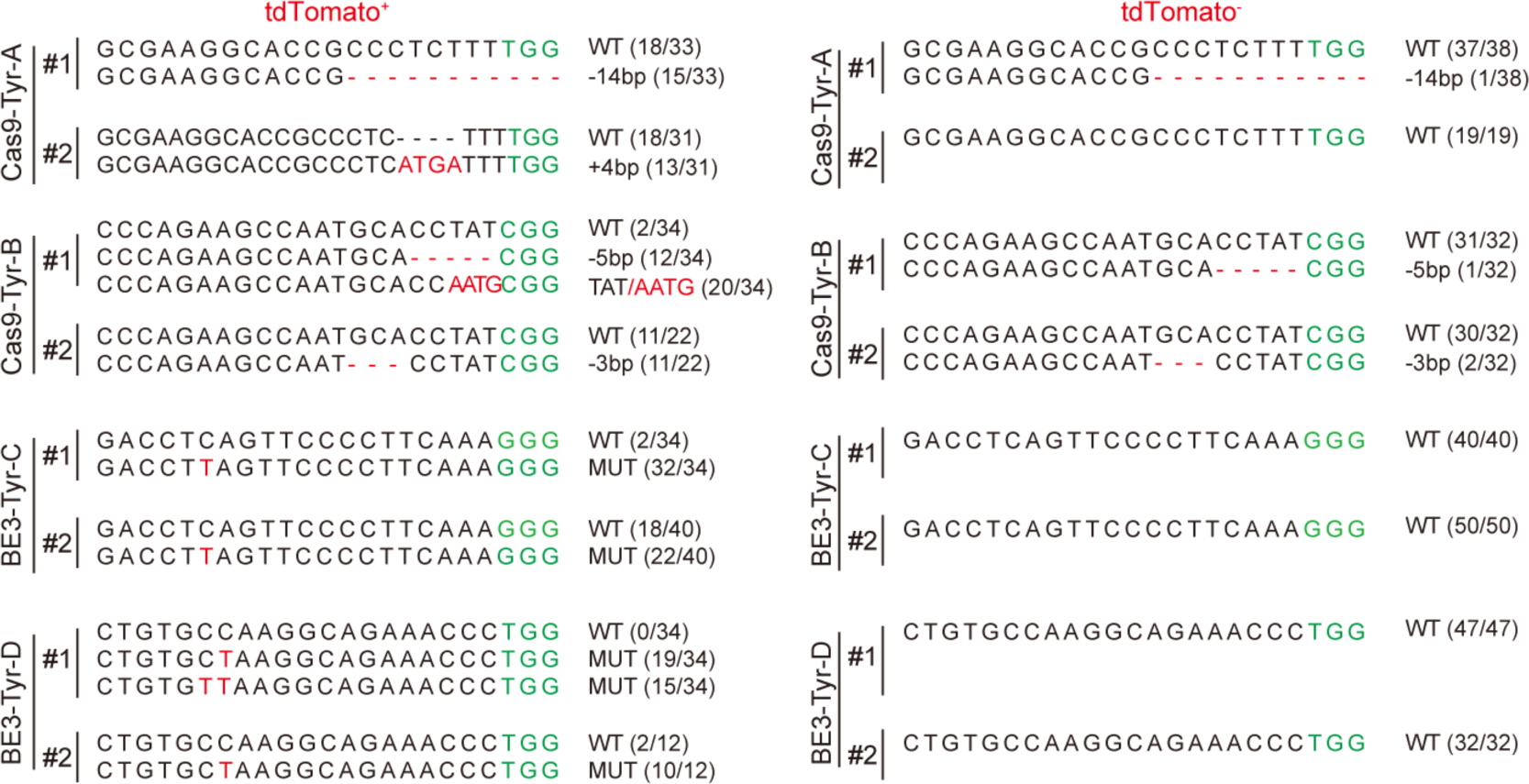
On-target sequences from WGS in tdTomato^+^ and tdTomato^-^ cells. WT and mutant sequences of the WGS results were shown as “WT” and “MUT”, the number before the slash shows the WT or mutant sequence reads, and the total reads were shown after the slash. Mutated sequences were marked in red color, and PAM was marked with green.

**Figure S5.**
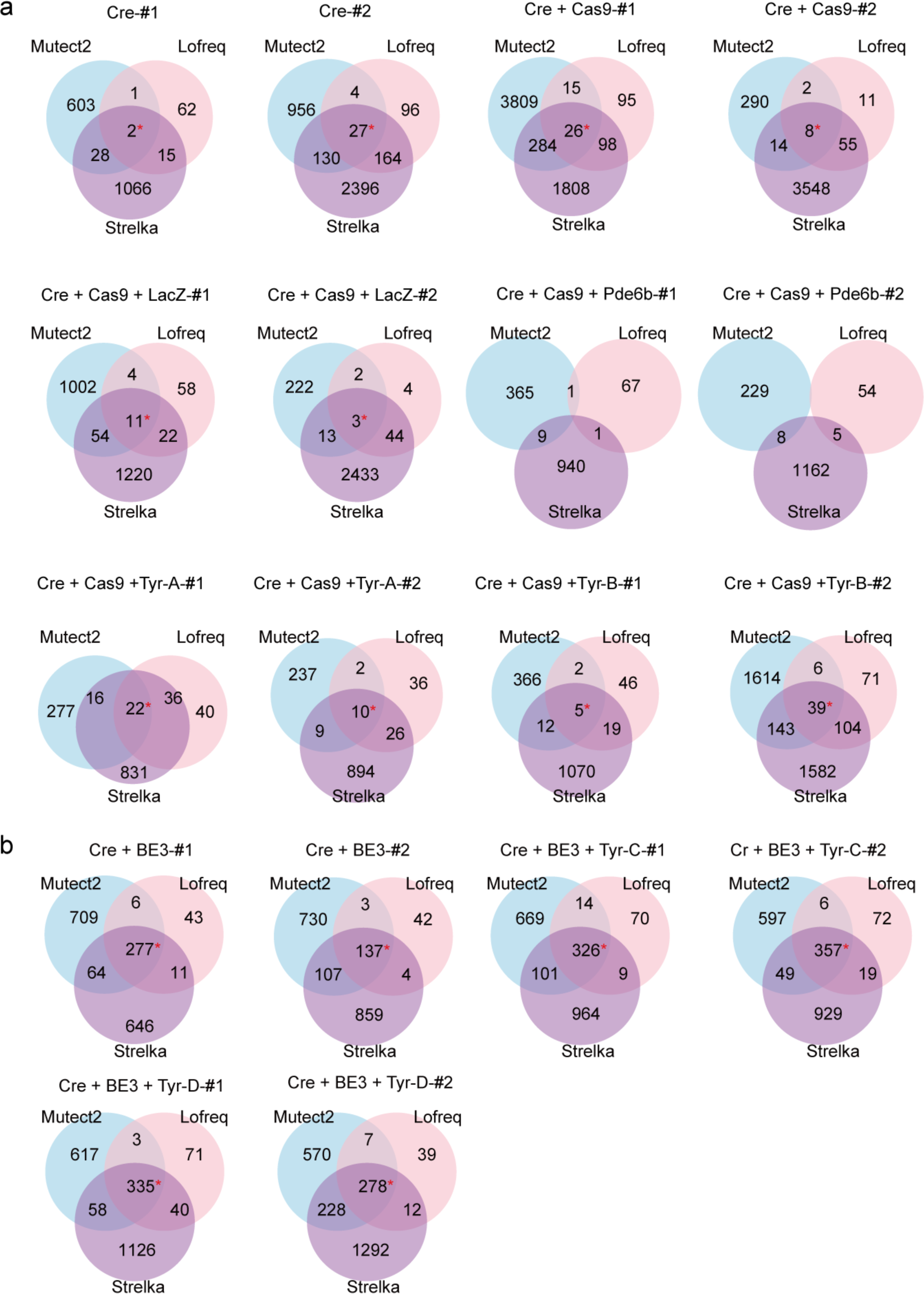
Venn diagrams of SNVs detected in each embryo by WGS data using the indicated software tools. **a**, SNVs detected in Cre or CRISPR/Cas9-treated samples. **b**, SNVs identified in BE3-treated embryos. ^*^The overlap SNVs with allele frequencies less than 10% were removed from the following analysis.

**Figure S6.**
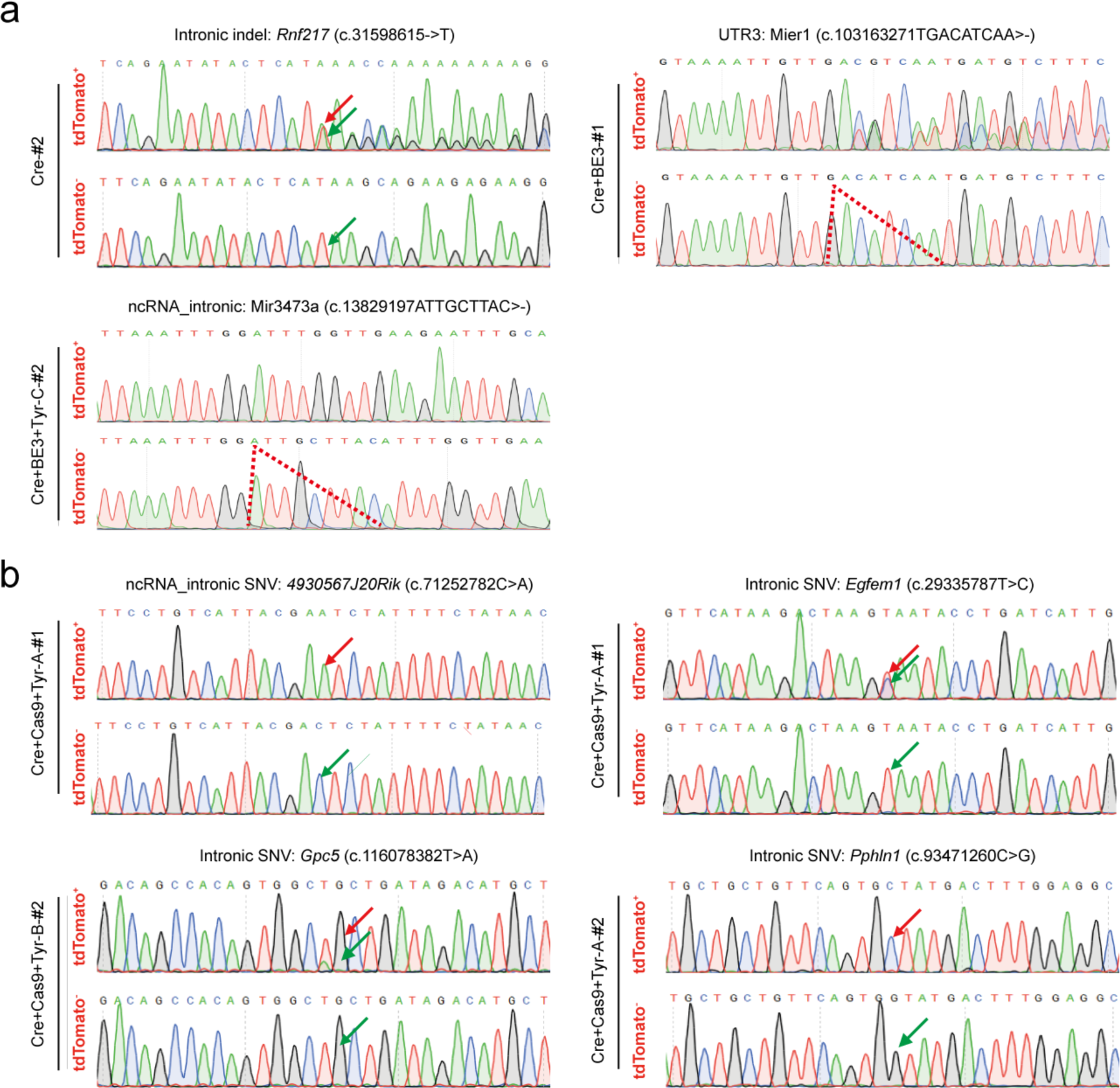
Representative Sanger sequencing chromatograms of mutations detected by WGS in Cre- or CRISPR/Cas9-treated embryos. **a**, Indels from indicated sample were PCR amplificated and Sanger sequenced. Green arrows, wild type; red arrows, inserted nucleotide. Red dot line, deleted nucleotides. **b**, SNVs from indicated samples were validated by Sanger sequencing. Green arrows, wild-type nucleotide; red arrows, mutated nucleotide. Primers were listed in table S16.

**Figure S7.**
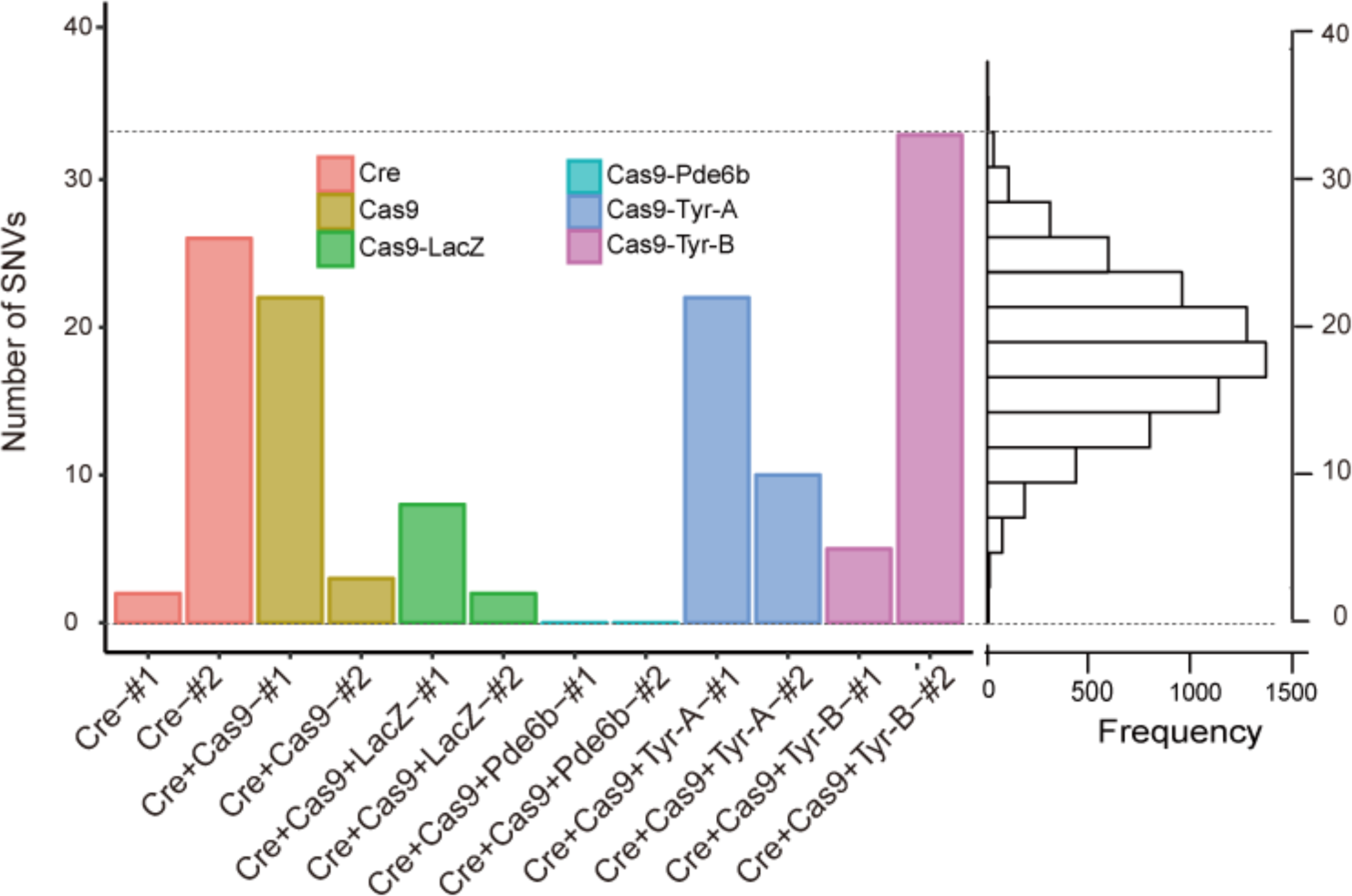
The number of SNVs detected from WGS data in embryos treated with Cre and CRISPR/Cas9. Embryos in the same group were indicated in the same color. The right panel is the distribution of simulated number of spontaneous mutations.

**Figure S8.**
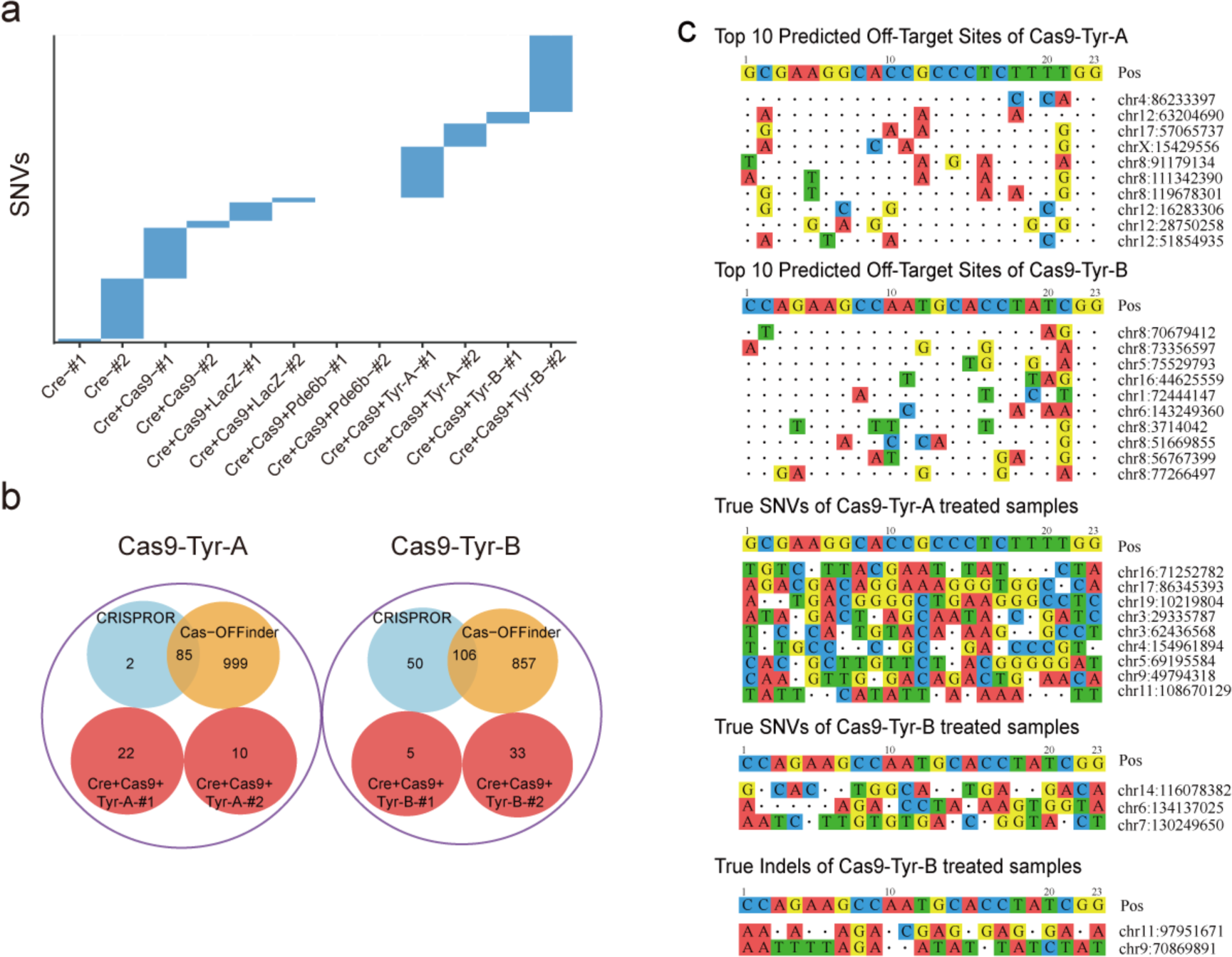
Off-target SNVs and indels identified from Cre- and CRISPR/Cas9-treated embryos. **a**, The SNVs identified from each embryo with Cre or CRISPR/Cas9 injected was mutually exclusive from the others. **b**, The overlap among SNVs detected from CRISPR/Cas9-treated embryos with predicted off-targets sites by Cas-OFFinder and CRISPOR. **c**, Sequence alignment of top 10 predicted off-target sites with Cas9-Tyr-A and Cas9-Tyr-B and the detected mutations from the WGS data with Cas9-Tyr-A and Cas9-Tyr-B.

**Figure S9.**
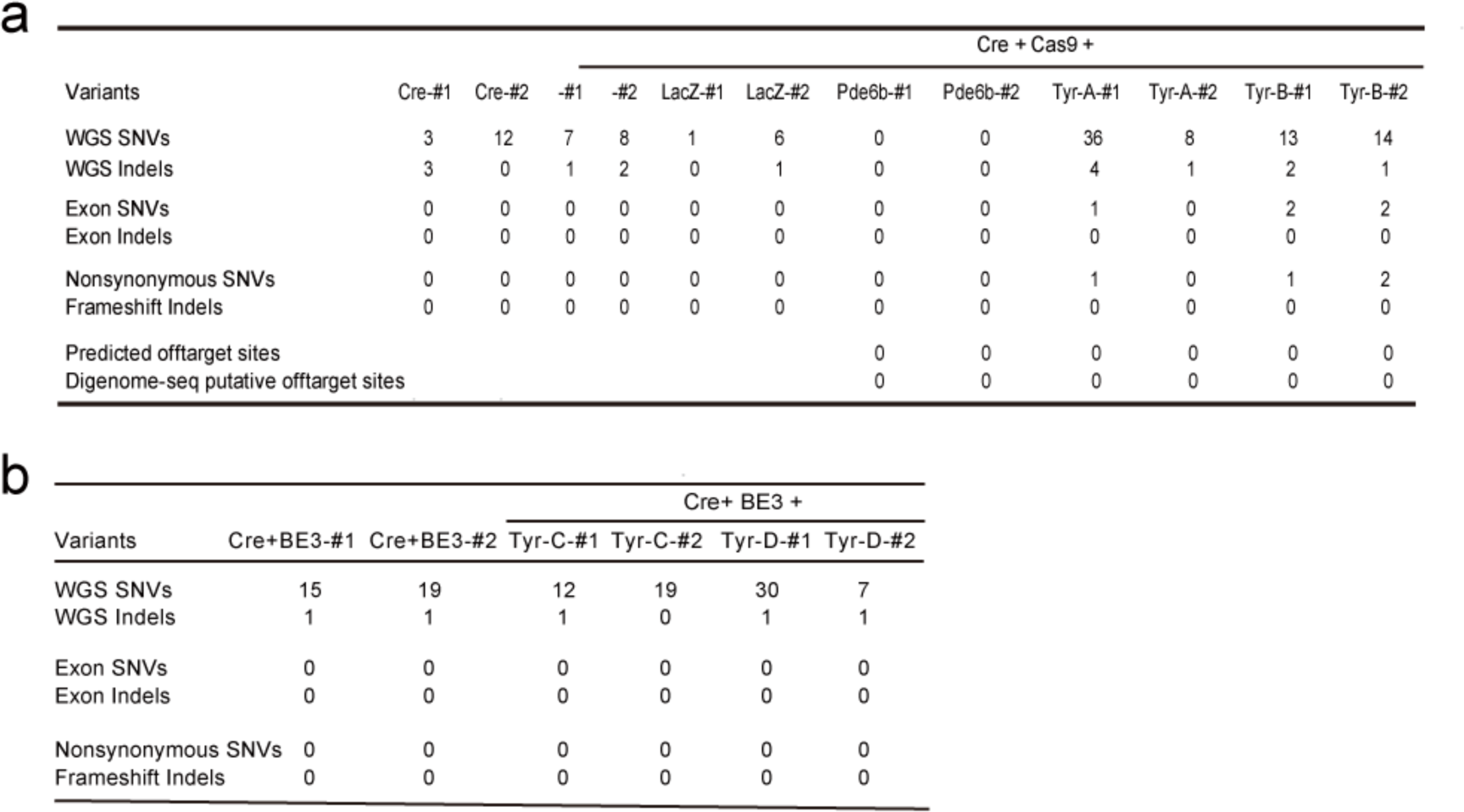
Summary of variants calling inversely from WGS data by comparing tdTomato^-^ with tdTomato^+^ cells from the same embryos. **a**, Converse variants calling from Cre- and CRISPR/Cas9-treated samples. **b**, Converse calling results from BE3-treated samples.

**Figure S10.**
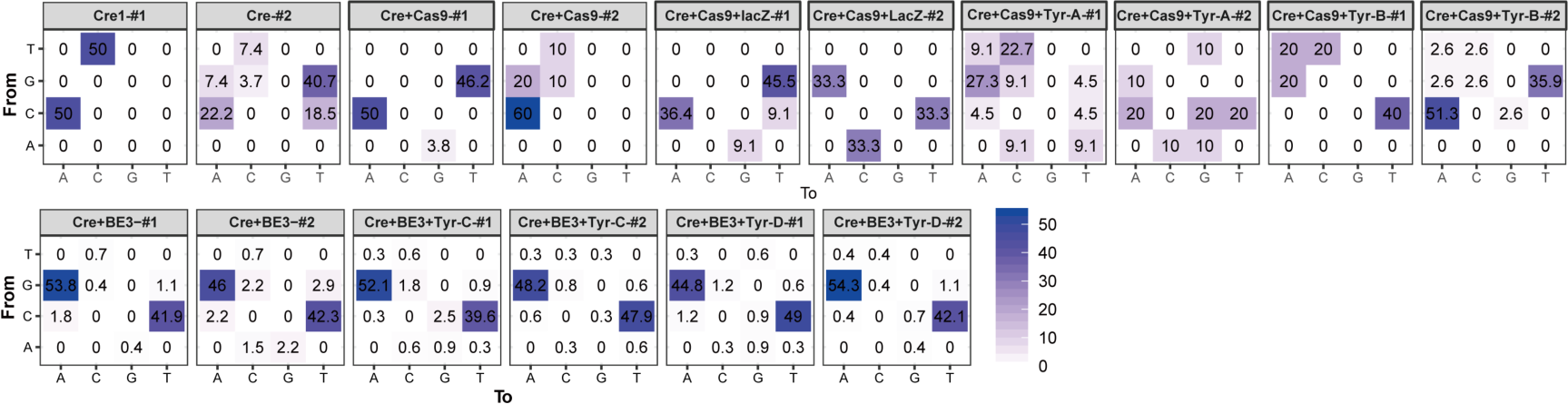
Mutation types of identified SNVs in each embryo included in the study. The number in each cell indicated the proportion of a certain mutation type, and deeper colors represented higher proportions.

**Figure S11.**
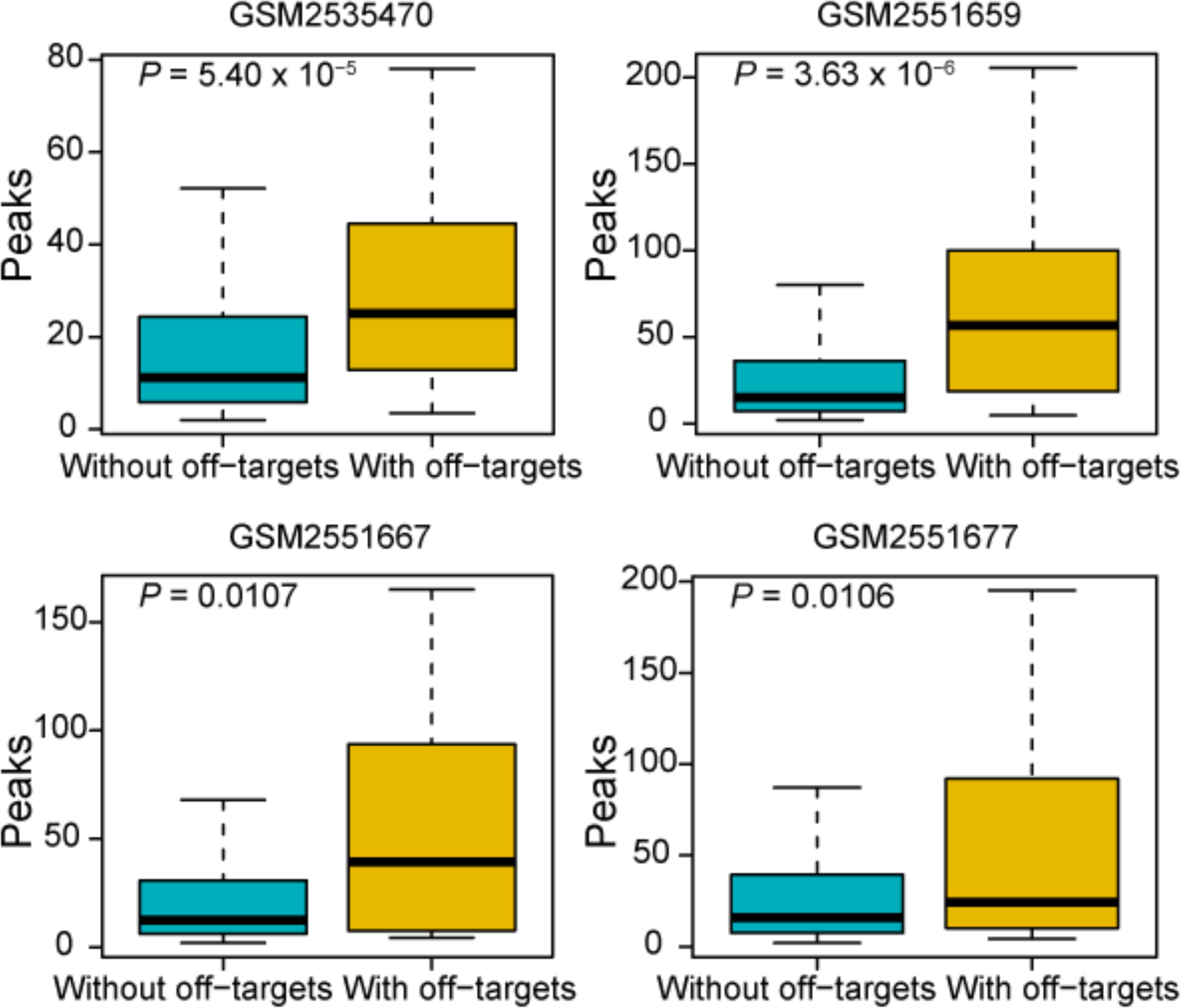
The comparison of peak regions with or without the identified off-targets from four datasets of Cistrome. The numbers on top of each boxplot indicated the GEO accession of applied datasets. *P*-values were calculated with Wilcoxon rank sum test.

**Figure S12.**
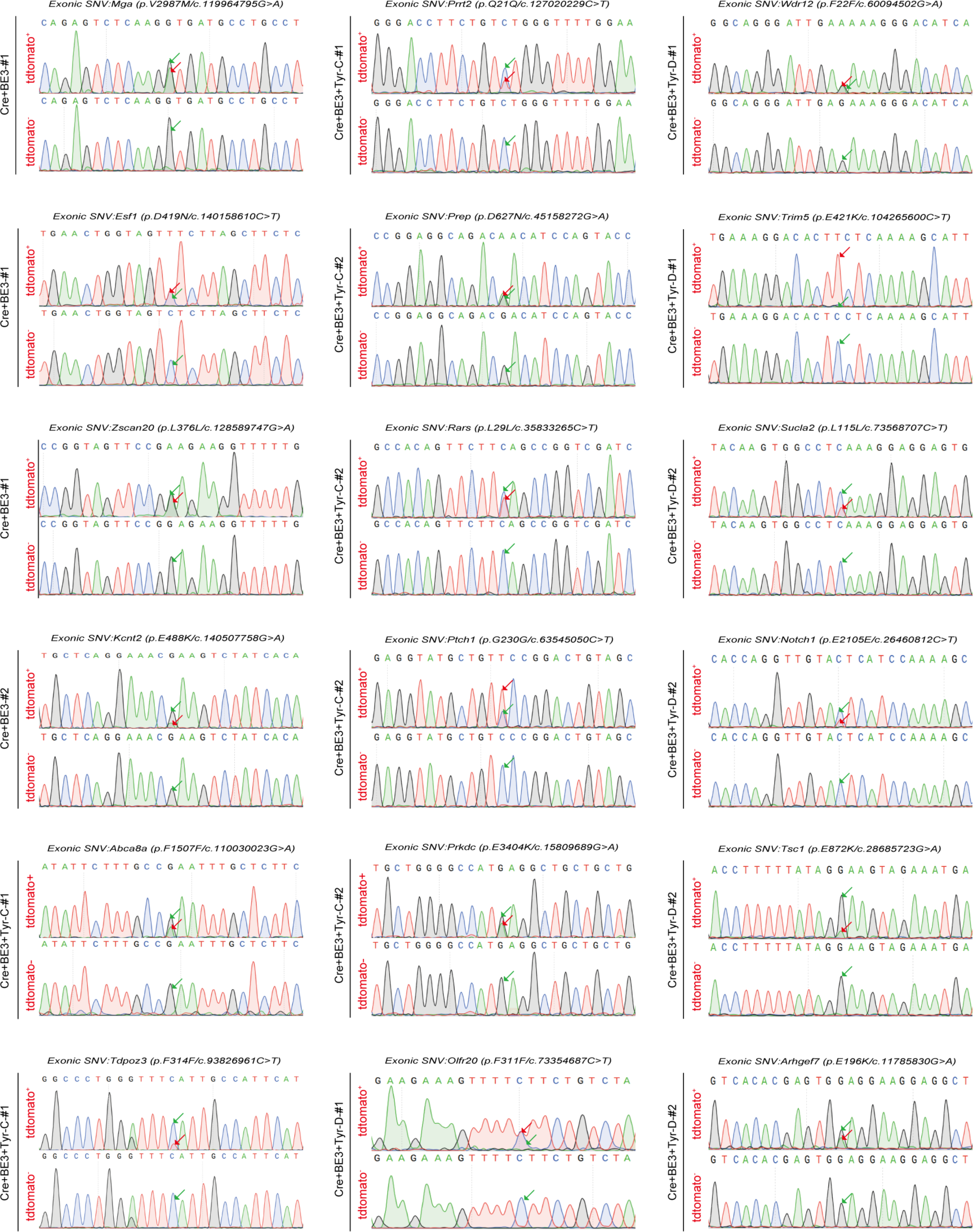
Sanger sequencing chromatograms of SNVs located on exons detected by WGS in BE3-treated embryos. SNVs from the indicated samples were validated by Sanger sequencing. Green arrows, wild-type nucleotide; red arrows, mutated nucleotide. Primers were listed in table S16.

## Matereials and methods

### Mice

Female C57BL/6 mice (4 weeks old) and heterozygous Ai9 (full name B6.Cg-Gt(ROSA)26Sortm9(CAG-td-Tomato)Hze/J; JAX strain 007909) male mice were used for embryo collection. ICR females were used for recipients. The use and care of animals complied with the guideline of the Biomedical Research Ethics Committee of Shanghai Institutes for Biological Science, Chinese Academy of Sciences.

### Generation of Cas9 mRNA, BE3 mRNA, Cre mRNA and sgRNA

T7 promoter was added to Cas9 coding region by PCR amplification of px260, using primer Cas9 F and R. T7-Cas9 PCR product was purified and used as the template for in vitro transcription (IVT) using mMESSAGE mMACHINE T7 ULTRA kit (Life Technologies). T7 promoter was added to sgRNA template by PCR amplification of px330. The T7-sgRNA PCR product was purified and used as the template for IVT using MEGA shortscript T7 kit (Life Technologies). T7 promoter was added to Cre template by PCR amplification. T7-Cre PCR product was purified and used as the template for *in vitro* transcription (IVT) using mMESSAGE mMACHINE T7 ULTRA kit (Life Technologies). Cas9 mRNA, Cre mRNA and sgRNAs were purified using MEGA clear kit (Life Technologies) and eluted in RNase-free water.

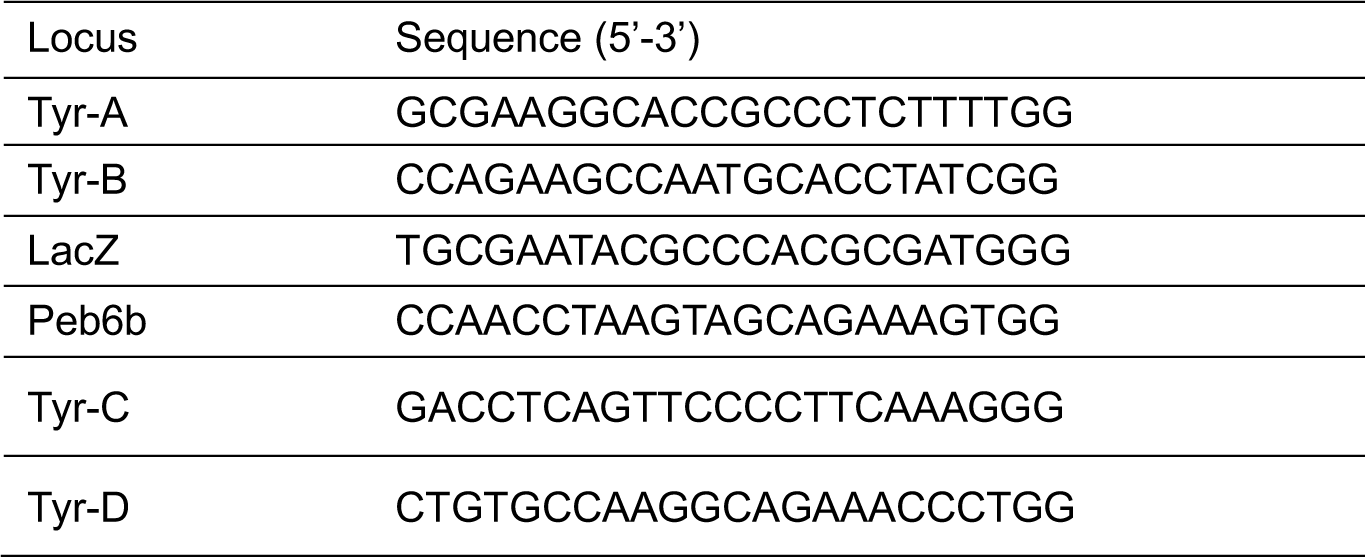
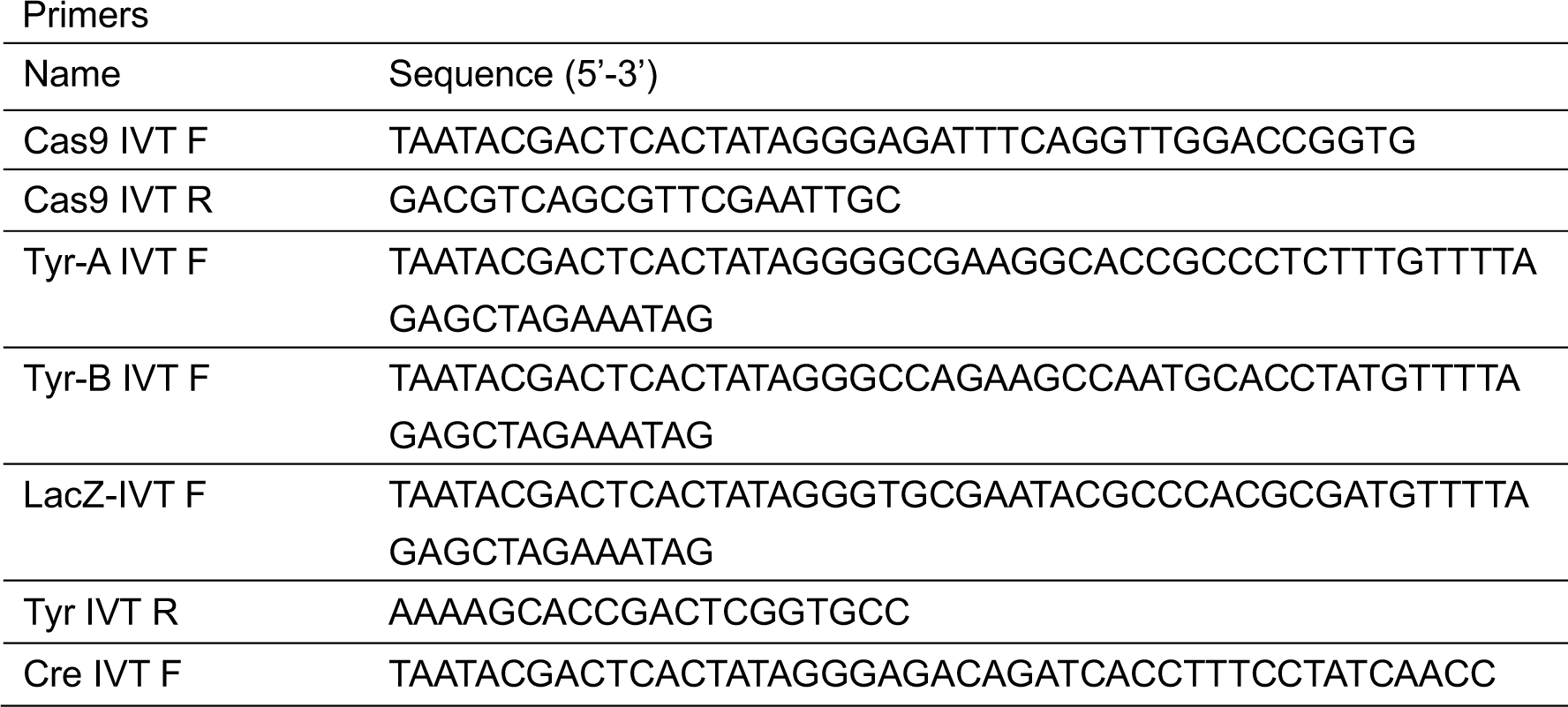
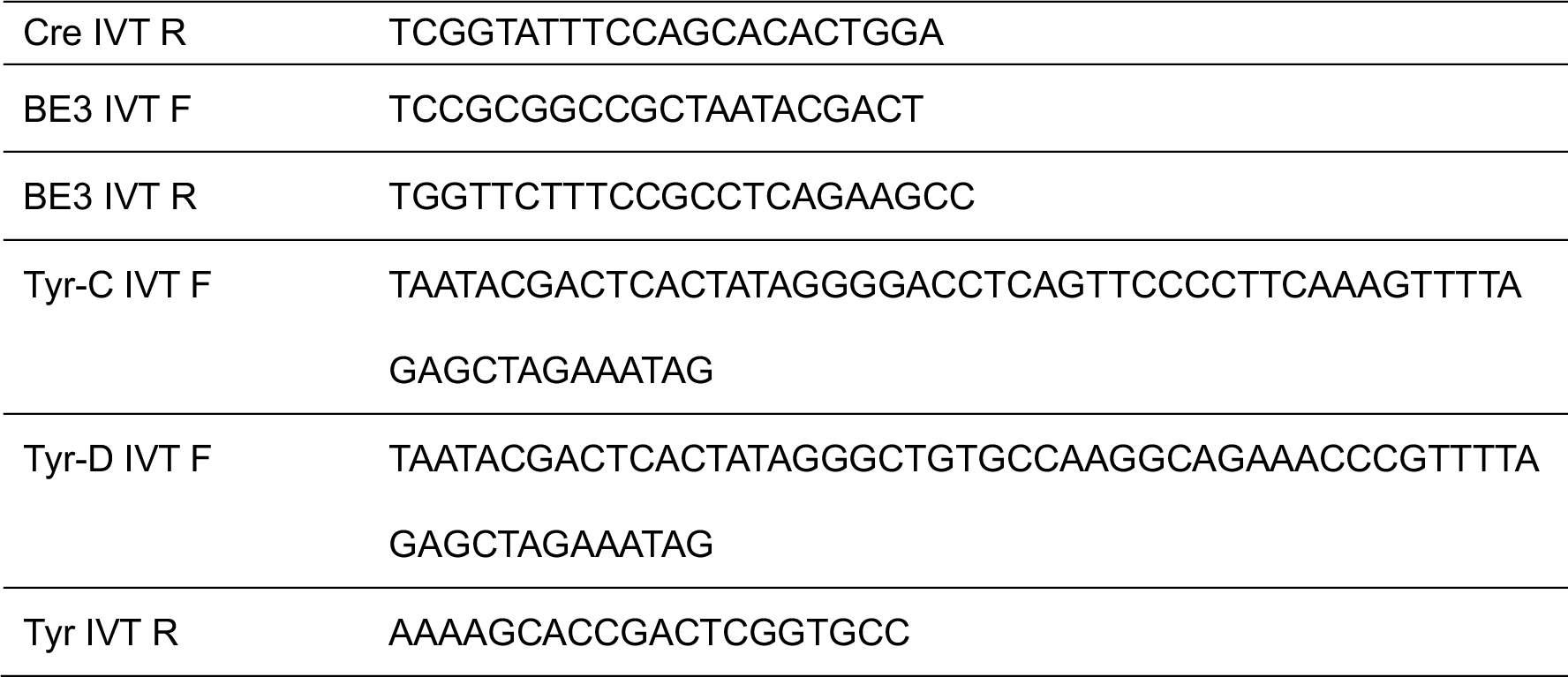

### 2-cell Embryo Injection, Embryo Culturing, and Embryo Transplantation

Super ovulated C57BL/6 females (4 weeks old) were mated to heterozygous Ai9 (full name B6.Cg-Gt(ROSA)26Sortm9(CAG-td-Tomato)Hze/J; JAX strain 007909) males, and fertilized embryos were collected from oviducts 23 h post hCG injection. For 2-cell editing, the mixture of Cas9 mRNA (50 ng/μl) or BE3 mRNA (50 ng/μl), sgRNA (50 ng/μl) and Cre mRNA (2 ng/μl) was injected into the cytoplasm of one blastomere of 2-cell embryo 48 h post hCG injection in a droplet of HEPES-CZB medium containing 5 μg/ml cytochalasin B (CB) using a FemtoJet microinjector (Eppendorf) with constant flow settings. The injected embryos cultured in KSOM medium with amino acids at 37°C under 5% CO2 in air for 2 hours and then transferred into oviducts of pseudopregnant ICR females at 0.5 dpc.

### Single-cell PCR Analysis

To pick up and transfer single cells, we used a glass capillary under a dissection microscope. Eight-cell mouse embryos were digested with acid Tyrode solution to remove the zona pellucida, and then the embryos were transferred into 0.25% trypsin and gently pipetted to separate individual blastomere. Finally, the blastomeres were washed into 0.25% trypsin for 7 to 10 times and transferred into a PCR tube. 1.5 μl lysis buffer containing 0.1% tween 20, 0.1% Triton X-100 and 4 μg/ml proteinase K was then pipetted into the tube. Each tube was centrifuged to facilitate the mix. The lysis was incubated at a temperature of 56°C for 30 min, followed by 95°C for 5 min. The products of the lysis program were used as templates in a nest PCR analysis. All the operations were operated carefully in order not to pollute the samples.

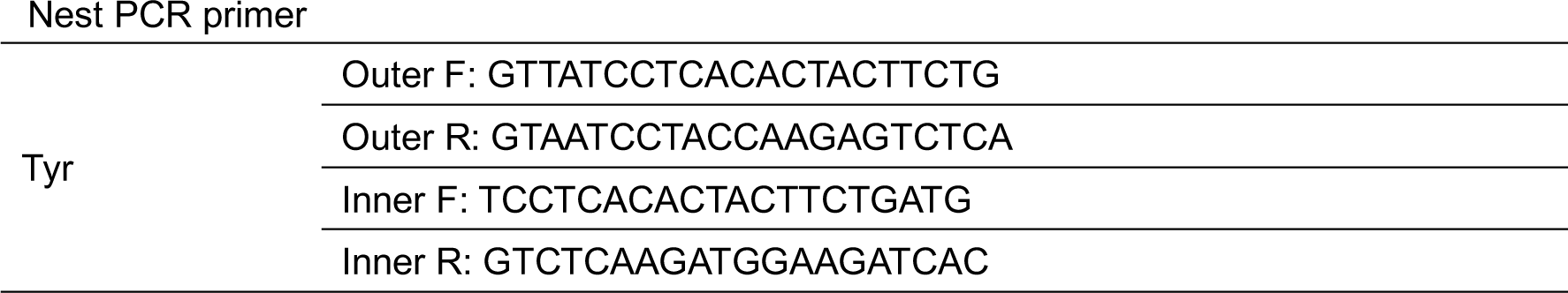

### TA Cloning and Genotyping

The PCR product was purified and ligated to pMD18-T vector and transformed to competent E. coli strain DH5α. After overnight culture at 37 °C, randomly selected clones were sequenced by the Sanger method. The genotypes of mutant E14.5 embryos were determined by PCR of genomic DNA extracted from cells. ExTaq was activated at 95°C for 3 min, and PCR was performed for 34 cycles at 95°C for 30 sec, 62°C for 30 sec, and 72°C for 1min, with a final extension at 72°C for 5 min. For blastocysts, after washing 6 times with KSOM, the single blastocyst was transferred directly into PCR tubes containing 1.5 μl embryo lysis buffer (0.1% tween 20, 0.1% Triton X-100 and 4 μg/ml proteinase K) and incubate for 30 min at 56°C, heat inactivate proteinase K at 95°C for 10 min. PCR amplification was performed using nest primer sets. ExTaq was activated at 95°C for 3 min, and PCR was performed for 34 cycles at 95°C for 30 sec, 62°C for 30 sec, and 72°C for 1min, with a final extension at 72°C for 5 min. Secondary PCR was performed using 0.5 μg primary PCR product and nested inner primer. PCR was carried out in the same reaction mixture. PCR product was gel purified and cloned using pMD-19t cloning kit (Takara) following the manufacturer’s instructions. Colonies were picked from each transformation and then Sanger sequencing to detect mutations. The primers used were:

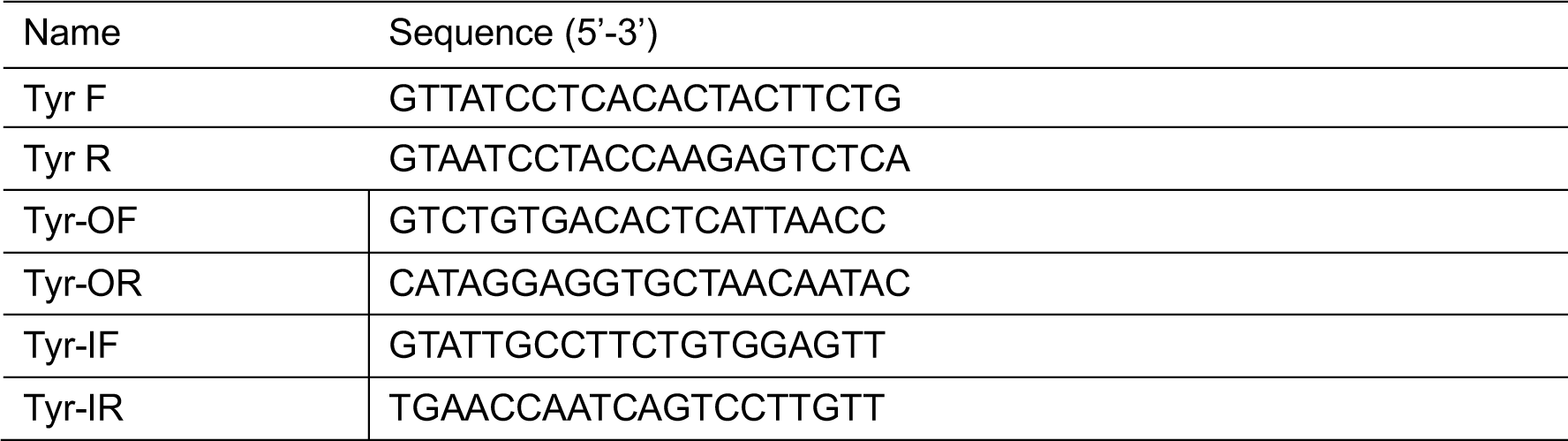

### FACS

To isolate cells, the prepared tissues were dissociated enzymatically in an incubation solution of 5 mL Trypsin-EDTA (0.05%) at 37°C for 30min. The digestion was stopped by adding 5 ml of DMEM medium with 10% Fetal Bovine Serum (FBS). Fetal tissues were then homogenized by passing 30-40 times through a 1ml pipette tips. The cell suspension was centrifuged for 6 min (800 rpm), and the pellet was resuspended in DMEM medium with 10% FBS. Finally, the cell suspension was filtered through a 40-μm cell strainer, and tdtomato^+^/tdtomato^-^ cells were isolated by FACS. Samples were found to be >95% pure when assessed with a second round of flow cytometry and fluorescence microscopy analysis.

### Whole genome sequencing and data analysis

Genomic DNA was extracted from cells by using the DNeasy blood and tissue kit (catalog number 69504, Qiagen) according to the manufacturer’s instructions. WGS was performed at mean coverages of 50x by Illumina HiSeq X Ten. BWA (v0.7.12) was used to map qualified sequencing reads to the reference genome (mm10). Picard tools (v2.3.0) was then applied to sort and mark duplicates of the mapped BAM files. To identify the genome wide *de novo* variants with high confidence, we conducted single nucleotide variation calling on three algorithms, Mutect2 (v3.5), Lofreq (v2.1.2) and Strelka (v2.7.1), separately(37-39). In parallel, Mutect2 (v3.5), Scalpel (v0.5.3) and Strelka (v2.7.1) were run individually for the detection of whole genome *de novo* indels(37, 39, 40). The overlap of three algorithms of SNVs or indels were considered as the true variants. We also marked variants overlapped with the UCSC repeat regions and microsatellite sequences or existed in the dbSNP (v138) and MGP (v3) databases. All the sequencing data were deposited in NCBI Sequence Read Archive (SRA) under project accession SRP119022.

To validate the on-target efficiency, we blasted the aligned BAM files with the on-target sites with e-value set as 0.0001. Potential off-targets of targeted sites were predicted using two previous reported algorithms, Cas-OFFinder (http://www.rgenome.net/cas-offinder/) and CRISPOR (http://crispor.tefor.net/) with all possible mismatches(21, 22).

The SNVs and indels were annotated with annovar (version 2016-02-01) using RefSeq database(41). Proto-oncogenes and tumor-suppressor genes were retrieved from the UniprotKB/Swiss-Prot database (2018_09). We downloaded five ATAC-seq bed peak files from the CistromeDB database with biological sources as embryonic and passing all quality controls *(28).* The five datasets retrieved included CistromeDB IDs “79877” (GSM2551659), “79976” (GSM2551677), “80493” (GSM2535470), “81049” (GSM2551664) and “81052” (GSM2551667). The off-target sites were mapped to the peak regions in each bed file based on the chromosome positions, and then the peaks regions with or without the off-targets were compared with each other by two-sided Wilcoxon rank sum test.

### Simulation of spontaneous mutations in the development of embryos

To estimate the number of spontaneous mutations from the 2-cell stage to E14.5 stage, the occurrence of single nucleotide mutations was simulated in silico considering the average sequencing coverage of 40 and allele frequency threshold of 10%. For each round of simulation, given the mutation rate of 1.8 x 10^-10^ and the size of mouse nuclear genome (2,785,490,220bp), we considered the replication process from the 2-cell stage to 16-cell stage, for mutations occurring after 16-cell stage will not be detected taken the allele frequency into consideration. Each cell may mutate or not during each replication, and once mutation occurs, the divided cells will inherit the mutation. Then the accumulated mutations and their wild-type alleles were randomly selected for sequencing with depth of 40, the selected mutations were summed as the number of spontaneous mutations for each round, and the same process was repeated for 10,000 times.

### In vitro cleavage of DNA

The genomic region flanking the sgRNA target site for each gene was PCR amplified and PCR amplicons was purified with the Universal DNA Purification Kit (Tiangen) according to the manufacturer’s instructions. Genomic DNA was purified from mice tail with the TIANamp Genomic DNA Kit (Tiangen) according to the manufacturer's instructions. The Cas9 ribonucleoproteins (1 μg) and sgRNA (1 μg) were pre-incubated at room temperature for 10 min to form RNP complexes. DNA (4 μg) was incubated with RNP complexes in a reaction buffer for 3 h at 37°C. Digested DNA was purified again with Universal DNA Purification Kit (Tiangen) after RNase A (100 μg/ml) was added to remove sgRNA.

### Digenome sequencing analysis

Libraries were subjected to WGS using Illumina HiSeq X Ten sequencer and the WGS was performed at a sequencing depth of 30x to 40x. Qualified reads were aligned to the mouse reference genome (mm10) by Isaac aligner with the following parameters: base quality cutoff, 15; keep duplicate reads, yes; variable read length support, yes; realign gaps, no. DNA cleavage sites were identified computationally using Digenome-seq2 (https://github.com/chizksh/digenome-toolkit2).

### Statistical analysis

R version 3.5.1 (http://www.R-proiect.org/) was used to conduct all the statistical analyses in this work. All tests conducted were two-sided, and the significant difference was considered at *P* < 0.05.

